# Quantitative imaging of corneal endothelial development reveals dynamic but resilient monolayer

**DOI:** 10.64898/2026.05.01.722310

**Authors:** R Ramarapu, WR Stoehr, M Miesen, N Amro, SM Thomasy, CD Rogers

## Abstract

The formation of functional corneal endothelial cells during development requires tight coordination between tissue-scale growth and cell-scale organization, yet how these processes are integrated in three dimensions remains poorly understood. Here, we combine high-resolution confocal imaging with quantitative analysis to reconstruct the morphogenesis of the chick corneal endothelium across embryonic development. Using a pipeline integrating 3D nuclear segmentation, Voronoi-based topological mapping, and spatial statistics, we link macroscopic globe expansion to single-cell geometry and lattice organization. We identify a multiphasic relationship between tissue growth and cell density, driven by temporal decoupling of organ expansion and proliferation. During early development, rapid globe expansion induces cellular stretching and spatial heterogeneity, followed by a phase of density accumulation and geometric refinement. Despite these dynamic conditions, the endothelial sheet maintains a robust monolayer architecture with minimal z-axis stratification. Quantitative topological analysis reveals that hexagonal packing is preserved from early stages and progressively refined through reduction of area variability and spatial clustering. Nearest-neighbor and Clark-Evans analyses demonstrate a transition from localized clustering to a more uniform spatial distribution, consistent with increasing packing regularity. Transient out-of-plane deviations coincide with key mechanical transitions, suggesting a role for 3D remodeling in accommodating mechanical stress. Concomitantly, junctional and cytoskeletal organization undergo progressive maturation. N-cadherin is established early at cell-cell interfaces, while Zonula Occludens-1 (ZO-1) transitions from diffuse localization to apically enriched tight junctions aligned with cortical actin. In parallel, microtubule organization becomes increasingly polarized to the apical domain, coinciding with the emergence of primary cilia. Together, these changes reflect coordinated establishment of epithelial polarity, barrier function, and mechanical stability. Overall, our study provides a multiscale, imaging-driven framework for understanding how epithelial tissues achieve and maintain geometric order under mechanical strain, establishing the corneal endothelium as an exemplar for linking developmental mechanics, 3D architecture, and epithelial topology.

**Summary Statement:** Using 3D imaging and quantitative analysis, this work reveals how corneal endothelial cells stay organized and form a regular pattern during growth, despite ongoing changes in tissue size and shape.

## Introduction

The formation of the vertebrate cornea requires precise coordination between macroscopic tissue expansion and cellular-scale architectural remodeling. The corneal endothelium, a posterior corneal monolayer that regulates stromal hydration and transparency, provides a tractable model for studying this process (Lwigale, 2015; Maurice, 1957). Corneal endothelial cells (CECs) originate from migratory, multipotent cranial neural crest (NC) cells, which invade the space between the lens and corneal epithelium during early embryogenesis (Cvekl and Tamm, 2004; Hay and Revel, 1969; Ibrahim et al., 2023; Steffek et al., 1979). Upon arrival, these mesenchymal cells undergo a mesenchymal-to-endothelial transition (MEndT) to form a functional barrier (Gage et al., 2005; Roy et al., 2015). This transition involves the loss of migratory mesenchymal characteristics, including irregular fibroblastic morphology, alongside the acquisition of apicobasal polarity, mature junctional complexes, and lateral contact inhibition (Babushkina and Lwigale, 2020; Campbell et al., 2017; Goldminz et al., 1979; Harrison et al., 2016; Petroll et al., 1999).

Between embryonic days 6 and 16, the avian corneal endothelium undergoes this structural transition while experiencing substantial, multidirectional mechanical strain associated with rapid globe expansion driven by intraocular pressure (Coulombre, 1957; Schmid, 2003). While the macroscopic growth dynamics of the developing eye are well characterized, the cellular mechanisms by which the endothelial lattice accommodates these forces remain poorly resolved (Lindner et al., 2017).

Recent advances in high-resolution imaging and computational analysis have revealed that developing corneal endothelia exhibit complex three-dimensional organization, dynamic cytoskeletal architecture, and transient topological rearrangements (Harrison et al., 2016; Ramarapu et al., 2026). However, longitudinal, quantitative reconstruction linking organ-scale growth to single-cell geometry and tissue topology is lacking. Here, we combine confocal imaging with a quantitative computational pipeline-including 3D nuclear segmentation, Voronoi-based topological mapping, and spatial statistical analysis-to reconstruct corneal endothelial morphogenesis across development. We show that temporal decoupling between globe expansion and cell proliferation drives a stretch-induced geometric equilibration, transforming a heterogeneous cranial NC cell population into a highly ordered hexagonal monolayer.

## Results

### Corneal endothelial cells exhibit multiphasic scaling relative to globe area expansion

To determine how endothelial cell density scales with organ growth, we quantified globe area and CEC nuclear density across embryonic days 6-16 (Fig. 1A, B, Table 1). Inter-rater reliability for globe measurements was high (Pearson R = 0.969, p < 0.001) (Fig. S1, Fig. S1). Globe area and nuclear density were strongly correlated (Pearson’s R = 0.92, p < 0.001) (Fig. 1B); however, analysis of temporal rates of change revealed a multiphasic relationship characterized by shifts in relative growth rates (Fig. 1C, Table S1). During early stages (days 6-9), globe area expanded rapidly (e.g., a 132% increase from 8.03 mm^2^ at day 6 to 18.80 mm^2^ at day 7, p < 0.001), while nuclear density remained relatively constant (2746 ± 731 cells/mm^2^; ns) (Fig. 1B, C). Between days 11 and 13, globe expansion slowed, reaching a transient plateau (day 11-12/12-13/13-14; ns), coinciding with significant increases in nuclear density (Fig. 1B, p < 0.01). After day 14, growth entered a steady phase in which gradual globe expansion was matched by stabilization of nuclear density, reaching a plateau at 5524 ± 410 cells/mm^2^ (Fig. 1B, C, Table S1).

**Table 1.** Globe metrics samples.

| Age | Embryos (N) | Globes (n) |
| --- | --- | --- |
| 6 | 7 | 14 |
| 7 | 5 | 10 |
| 8 | 4 | 8 |
| 9 | 4 | 8 |
| 10 | 4 | 8 |
| 11 | 5 | 10 |
| 12 | 8 | 16 |
| 13 | 11 | 20 |
| 14 | 7 | 13 |
| 15 | 8 | 15 |
| 16 | 10 | 20 |

**Figure 1.**
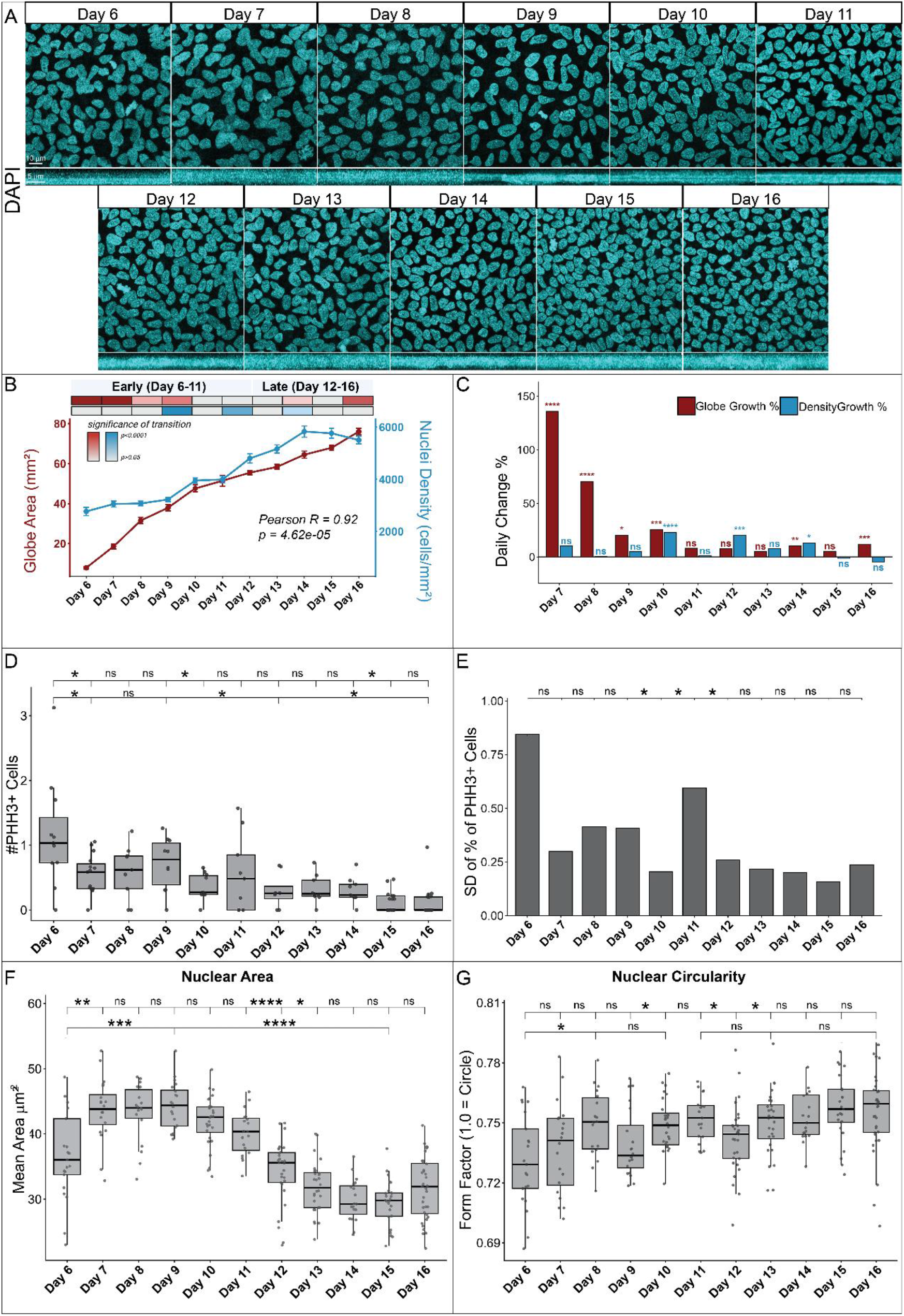
Relationship between globe area and nuclear density. A) Maximum intensity projections of DAPI demonstrates developing CEC nuclear distribution and overlap. Transverse MIPs in bottom panel. (B) Graph shows globe area (mm^2^) maintains a steady growth from Day 6 to Day 11, followed by a plateau together with CEC nuclear counts from Day 6 to Day 16 demonstrates stage-dependent increases in cell density. Significance of stage-wise transitions are represented by colored intensity boxes. Ocular measurements obtained using ImageJ. (C) Bar chart demonstrating % daily change values for globe area (red) and nuclear density (blue) (D, E) Graph shows number of PHH3-positive cells across developmental time and standard deviation across sample days. (F) Nuclear area demonstrating an oscillating pattern across development as globe size increases from day 6-9 following with a gradual reduction. (G) Quantification of nuclear circularity from Day 6-D16 shows significant reductions in circularity between days 9-10 and 11-12. Circularity trend reliably increases across developmental days 6-8. For N, refer to Table 1,2. Scale bars are as shown.

To investigate the basis of these density changes, we assessed proliferation and morphometric parameters. DAPI/ phosphorylated histone H3 (PHH3) co-staining showed that density increases were not driven by a discrete proliferative burst. Instead, the PHH3 index was highest on day 6 (1.03%) and declined sharply by day 7 (0.58%, p < 0.001), with further reduction between days 9 and 10 (p < 0.05) (Fig. 1D, E). Thereafter, proliferation remained at a low baseline (~0.5-1.0%) until a further decline at days 14-16 (p < 0.05) (Fig. 1D, E). Variance analysis (Fligner-Killeen) revealed that early baseline proliferation was spatially heterogeneous. Between days 9 and 11, PHH3 variance fluctuated significantly (p < 0.05), indicating localized and unsynchronized divisions. As the tissue matured into a more permanent lattice, this variance decreased, suggesting more uniform proliferative activity (Fig. 1D, E).

**Table 2.** Nuclear and PHH3 Segmentation Samples.

| ***Nuclear Segmentation*** |  |  |
| --- | --- | --- |
| Age | Embryos (N) | Images (n) |
| 6 | 13 | 21 |
| 7 | 10 | 20 |
| 8 | 9 | 18 |
| 9 | 10 | 20 |
| 10 | 12 | 28 |
| 11 | 11 | 19 |
| 12 | 13 | 29 |
| 13 | 15 | 28 |
| 14 | 9 | 19 |
| 15 | 13 | 25 |
| 16 | 15 | 34 |
| ***PHH3 Segmentation*** |  |  |
| 6 | 3 | 11 |
| 7 | 3 | 15 |
| 8 | 3 | 9 |
| 9 | 3 | 10 |
| 10 | 3 | 10 |
| 11 | 3 | 9 |
| 12 | 3 | 8 |
| 13 | 3 | 9 |
| 14 | 3 | 9 |
| 15 | 3 | 18 |
| 16 | 3 | 18 |

Morphometric analysis indicated that early density dilution was accommodated by cell hypertrophy: nuclear area increased significantly from day 6 to a peak at day 9 (Fig. 1F). Following this, as globe growth slowed, continued baseline proliferation led to tissue compaction and a sustained decrease in nuclear area from day 9 to day 15 (Fig. 1F). Nuclear circularity increased from day 6 through 8 but remained relatively stable overall excluding significant reductions on days 9 and 12 (Fig. 1G), corresponding to phases of maximal stretching and compaction, respectively.

### Progressive geometric equilibration and stabilization of the corneal endothelial lattice

To quantify tissue topology, we generated Voronoi tessellations from nuclear centroid coordinates (Fig. 2A). Spatial mapping revealed progressive refinement of the endothelial lattice toward a regular hexagonal organization (Fig. 2B). Analysis of a comprehensive geometric trajectory revealed a resilient hexagonal motif (Fig. 2C). Contrary to models of early developmental chaos predicted in more mesenchymal tissues, hexagons represented the highest relative proportion of cells across all observed stages. However, the lattice underwent significant refinement as the tissue matured (Fig. 2C). During the early expansion phase (day 6-9), the proportion of hexagons rose significantly from day 6 through 8, following which it maintained a stable plateau of ~42% (Fig. 2D). This topological refinement was accompanied by a reduction in intercellular area variability. The coefficient of variation (CV) of Voronoi polygon areas was highest on day 6 (~30.55%) and decreased significantly by day 9 (~20.77%, p < 0.001), followed by a further gradual decline to ~14.93% by day 16 (Fig. 2E). These results indicate progressive homogenization of cell packing.

**Figure 2.**
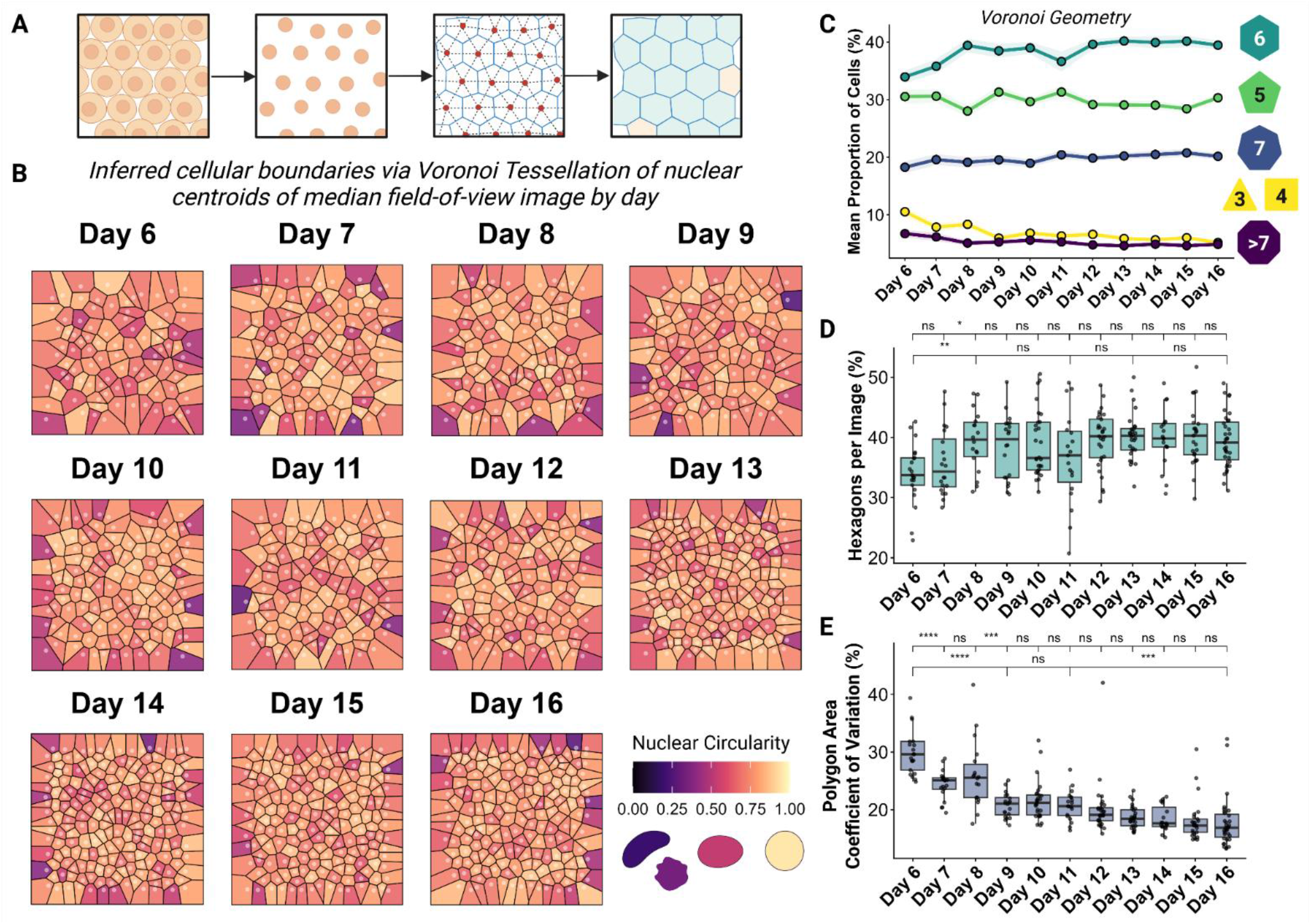
Progressive geometric equilibration of the developing corneal endothelial lattice. (A) Schematic illustration of Voronoi tessellations derived from 2D nuclear centroid coordinates, utilized to mathematically model the topological architecture of the monolayer. (B) Representative topological maps of the CEC monolayer from embryonic day 6 to 16. Voronoi polygons are colored by nuclear circularity. (C) Line plot detailing the relative frequency distribution of polygon geometries across developmental stages. (D) Box plot quantifying the hexagonality as a percentage of polygonality. (E) Box plot illustrating intercellular area variance, quantified as the coefficient of variation of Voronoi polygon areas. Significance assessed via Wilcoxon rank-sum testing; *p < 0.05, **p < 0.01, ***p < 0.001, ****p < 0.0001. For N, refer to Table 2.

### Spatial clustering resolves into a regularized distribution

To further characterize spatial organization, we analyzed nuclear nearest-neighbor distance (NND). Mean NND remained stable between days 6 and 9 but decreased significantly after day 9 as nuclear density increased, stabilizing by days 15-16 (Fig. 3A). The standard deviation (SD) of NND, reflecting spatial heterogeneity, was highest on day 6 and decreased progressively, with a marked drop on day 10 (Fig. 3B). This indicates increasing spatial uniformity.

**Figure 3.**
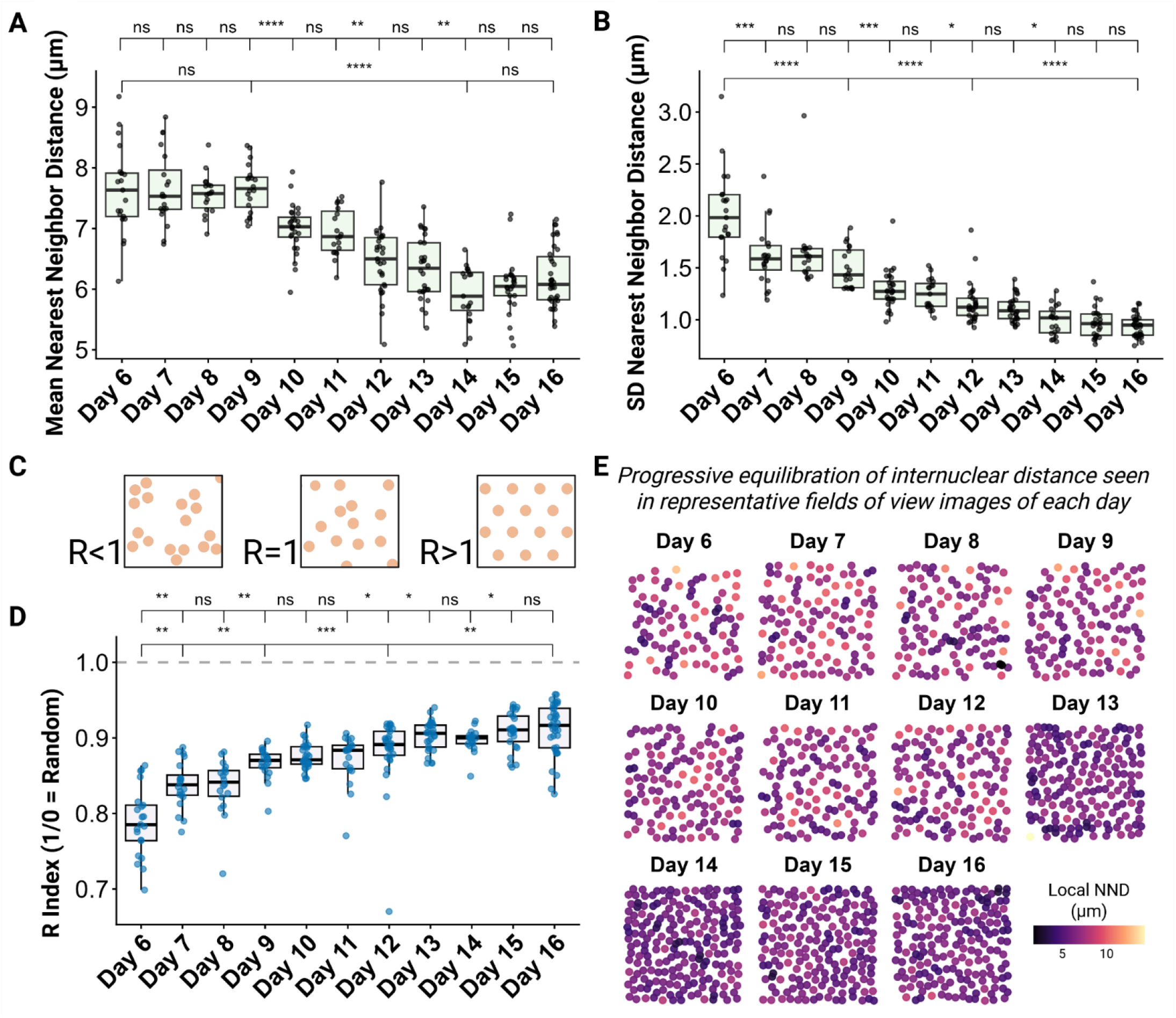
Inter-nuclear spacing dynamics reveal transition from localized clustering to regularized random lattice repulsion. (A) Boxplot quantifying the average nearest neighbor distance (NND) across developmental stages. (B) Boxplot of standard deviation (SD) of the NND, representing spatial variance. (C) Schematic illustrating Clark-Evans Aggregation Index. (D) Boxplot demonstrating Clark-Evans R index across developmental stages. (E) Representative 2D spatial coordinate maps of CEC nuclei, colored by local NND. Significance assessed via Wilcoxon rank-sum. *p < 0.05, **p < 0.01, ***p<0.001, ****p<0.0001. For N, refer to Table 2.

To determine the degree of underlying mathematical order driving this spacing, we next calculated the Clark-Evans aggregation index (R). The tissue exhibited a significant increase from R = 0.78 ± 0.04 at day 6 to ~0.91 on day 16 (p < 0.0001), indicating a transition from clustered to more uniformly distributed cell positions (Fig. 3C, D). Although R remained below 1, consistent with biological variability, this shift reflects increasing spatial regularization. This transition was visually confirmed through spatial coordinate mapping colored by local NND where early stages showed localized clustering, which progressively resolved into a dense and spatially uniform monolayer by later stages (Fig. 3E).

### The corneal endothelial sheet maintains monolayer organization in 3D

To assess the three-dimensional organization of the CECs over developmental time, we quantified nuclear deviation from the tissue midplane (ΔZ) using 3D spatial regression of CEC nuclei. Topological mapping of the monolayer colored by local Z-axis deviation (Fig. 4A) revealed distinct structural phases. During the peak PHH3 index on day 6, the monolayer exhibited localized, focal peaks of z-axis stacking, visually appearing as highly uneven terrain (Fig. 4A-C’, Fig. S2, S3). By day 7, correlating with the sharp drop in PHH3 index, the unevenness of the topological plane was reduced. The mean Median Absolute Deviation (Z-MAD) from independent embryos per stage was visualized using a line graph (Fig. 4B). The trend aligned with the visual maps as early topological variance is seen on day 6, which drops by day 7 to establish a generally low baseline. The data additionally revealed a reproducible, mild topological elevation on day 12, visible in both the quantitative Z-MAD trace, localized topological maps, and 3D projection of nuclei surfaces (Fig. 4A, B, D, D’). Linear mixed-effects modeling detected no significant global changes in Z-MAD across stages (p > 0.05), indicating that the endothelium maintains a stable monolayer architecture despite dynamic remodeling (Fig. S3). Rather than indicating a lack of biological activity, this lack of global variance likely confirms the CEC maintains a highly resilient, monolayer architecture, gaining more order through its developmental expansion.

**Figure 4.**
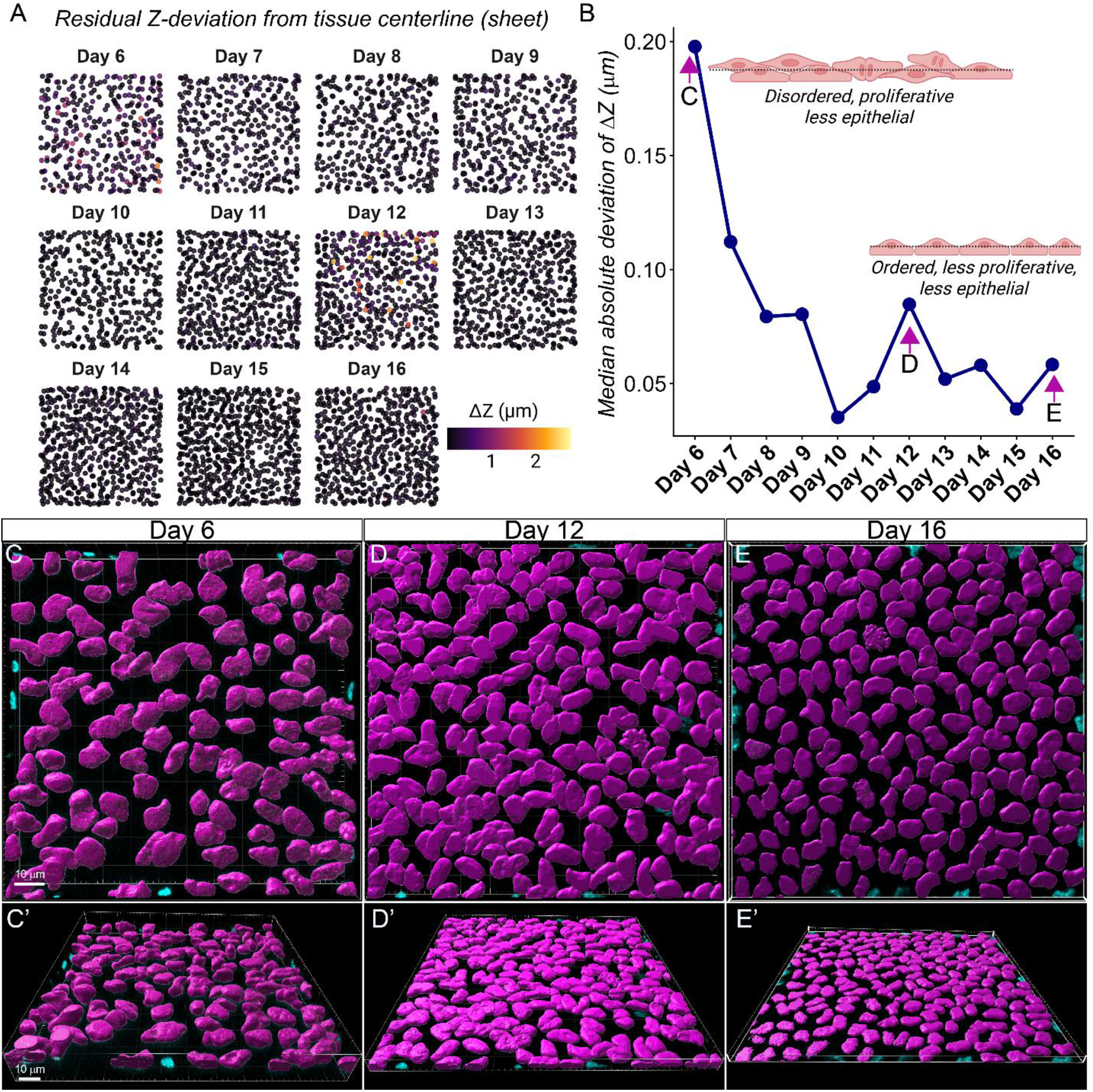
Endothelial sheet maintains relatively strict monolayer identity while accommodating transient topological perturbations. (A) Representative 3D topological maps displaying nuclear seed positions within a specified field of view, colored by their absolute residual deviation from the theoretical tissue center plane (ΔZ). (B) Line graph quantifying the mean absolute deviation of the z-plane (Z-MAD) across developmental time points. Linear mixed-effects modelling (LMM) over N = 3 biological replicates per stage confirmed no global, statistically significant stratification across the developmental timeline (p_LMM_ >0.05, 341-553 cells 3D segmented per sample range). 3D volumetric reconstruction of CEC nuclei at days (C, C’) 6, (D, D’) 12, and (E, E’) 16. For N, refer to Table 2.

### Establishment of corneal endothelial junctional complexes

Given that our analysis indicates the CEC forms a continuous monolayer by embryonic day 6, we next examined the establishment and maturation of apical junctional complexes that stabilize the tissue prior to terminal differentiation and pump function. To visualize junctional organization, we performed immunohistochemistry for N-cadherin (NCAD), together with filamentous actin (F-actin) and DAPI, and imaged flat-mounted corneas from the apical surface.

As early as day 6, NCAD was detectable at cell-cell interfaces, where it localized to cell membranes, and this localization was maintained through day 11 (Fig. 5A-F). Orthogonal (transverse) projections revealed progressive apical enrichment and sharpening of both NCAD and F-actin signals over time (Fig. 5A-F, bottom panels), indicating gradual maturation and refinement of junctional architecture as the endothelial lattice becomes more ordered. Consistent with our quantitative analysis of nuclear organization (Fig. 2), 3D nuclear surface reconstructions incorporating NCAD and F-actin showed that cells were relatively dispersed between days 6 and 10, followed by a progressive decrease in cell and nuclear size and increased spatial regularity by day 11 (Fig. 5G).

**Figure 5.**
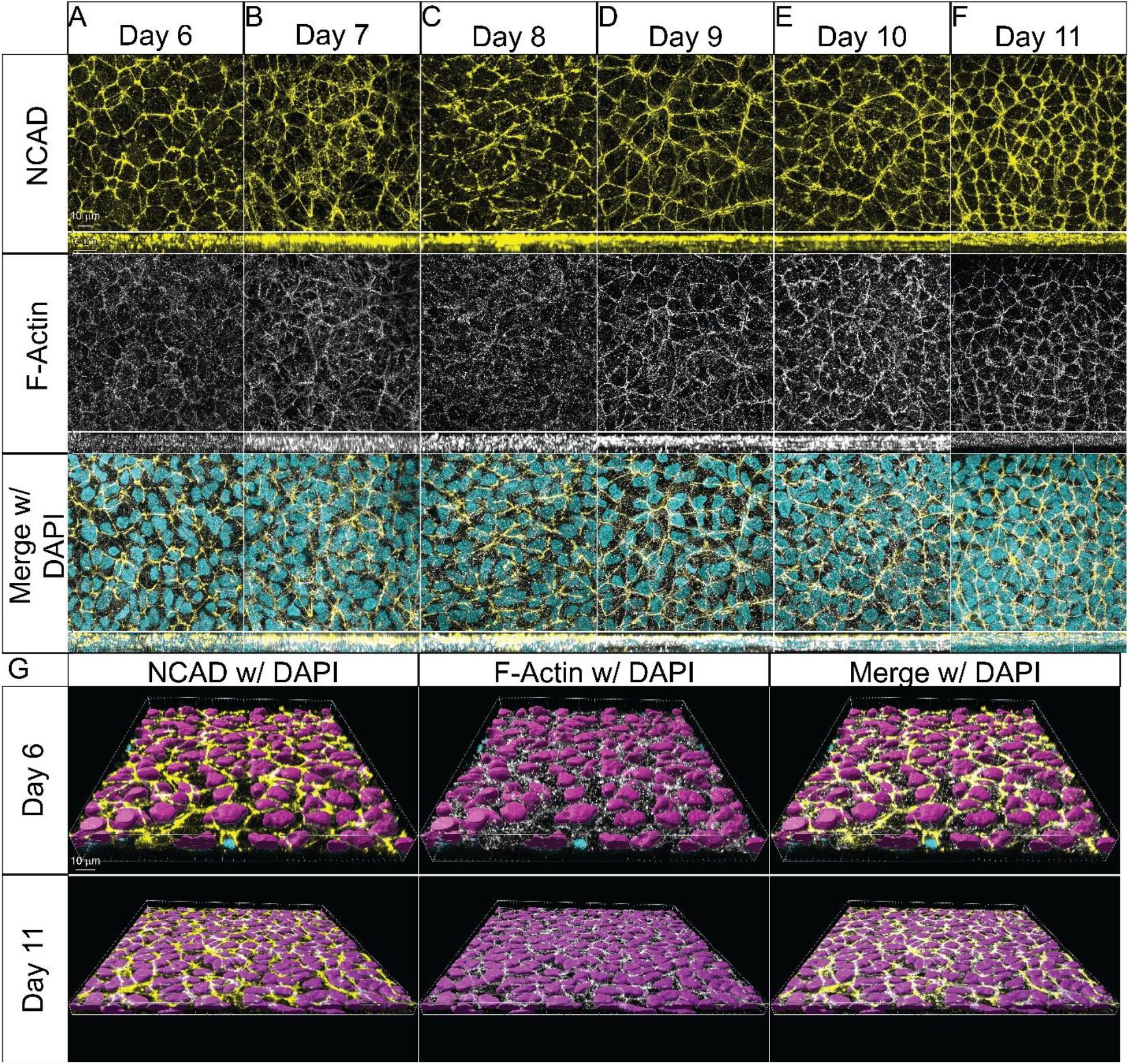
Progressive establishment and apical refinement of corneal endothelial adherens junctional complexes. (A-F) Representative confocal images of flat-mounted chick corneal endothelium from embryonic days 6-11 after IHC for N-cadherin (NCAD, adherens junctions), and stain for F-actin, and DAPI (nuclei), imaged from the apical surface. NCAD localizes to cell-cell interfaces from day 6 and is maintained throughout development. Bottom panels show corresponding orthogonal (transverse) projections highlighting apical-basal distribution of NCAD and F-actin signals. Progressive apical enrichment and sharpening of junctional signals are observed over time. (G) Representative 3D nuclear reconstructions integrating NCAD and F-actin. Early stages (days 6-10) show relatively dispersed cells with larger nuclear profiles, whereas later stages (day 11) exhibit reduced cell and nuclear size and increased spatial regularity, consistent with lattice refinement. Scale bars: flatmount and 3D-10 μm, transverse insets-5 μm.

While adherens junctions maintain cell-cell adhesion in CECs, tight junctions are critical for barrier function, and therefore we visualized maturation of tight junctions via ZO-1 IHC. ZO-1 signal was diffuse across the apicobasal plane and cytosol within the cells at day 11, but by day 12, the signal became membrane localized and overlapped with the cortical F-actin signal as the apical cell boundaries were refined (Fig. 6A-F). By day 16 both the ZO-1 and F-actin signals mirrored the hexagonal-dominant patterning seen in the nuclear positioning and NCAD and analyses (Fig. 2, 5). Orthogonal projections revealed similar progressive apical enrichment and sharpening of both ZO-1 and F-actin signals between days 11 to 16 (Fig. 6A-F, bottom panels), indicating gradual maturation and refinement of junctional architecture as the endothelial lattice becomes more ordered. 3D reconstructions incorporating ZO-1, F-actin, and nuclear signals showed a progressive refinement of ZO-1 signal to intercellular tight junctions (Fig. 6G).

**Figure 6.**
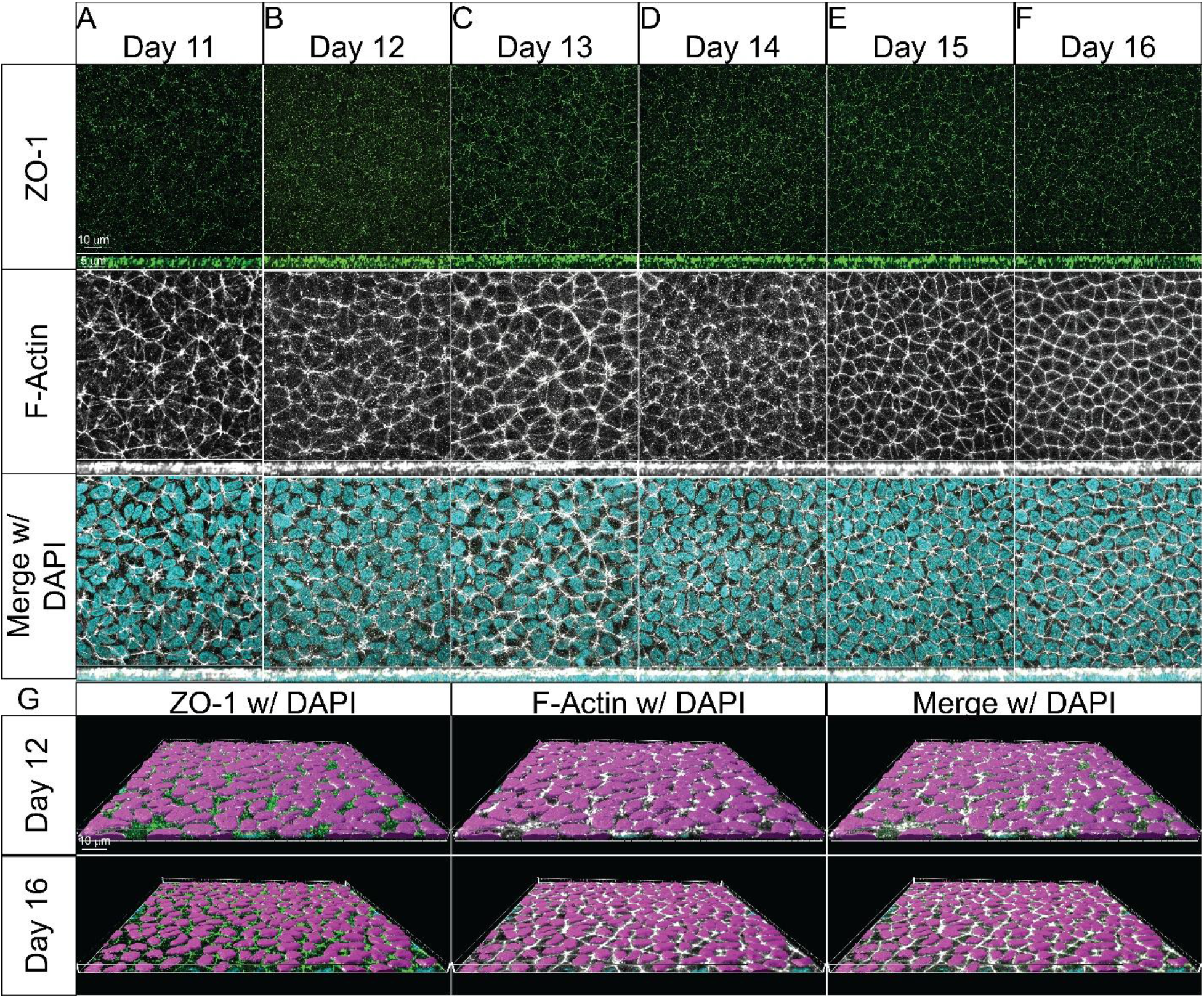
Maturation and apical refinement of tight junctions during corneal endothelial development. (A-F) Representative confocal images of flat-mounted chick corneal endothelium from embryonic days 11-16 after IHC for ZO-1 (tight junctions), F-actin, and DAPI (nuclei), imaged from the apical surface. At day 11, ZO-1 signal is diffuse across the apicobasal axis and within the cytosol. By day 12, ZO-1 becomes progressively enriched at cell–cell interfaces, co-localizing with cortical F-actin as cell shape becomes more regular. By day 16, both ZO-1 and F-actin display a clear hexagonal pattern consistent with mature endothelial organization. Bottom panels show corresponding orthogonal (transverse) projections, highlighting progressive apical enrichment and sharpening of junctional signals over time. (G) Representative 3D nuclear reconstructions integrating ZO-1 and F-actin, illustrating the progressive restriction of ZO-1 to intercellular tight junctions and increased spatial organization of the endothelial lattice. Scale bars: flatmount and 3D-10 μm, transverse insets-5 μm.

### Cytoskeletal architecture is progressively polarized during corneal endothelial development

Our analyses of nuclear organization and junctional maturation indicate that the corneal endothelium transitions from a relatively disordered sheet to a highly organized monolayer with refined apical junctions. To examine how cytoskeletal organization contributes to this process, we performed IHC for α-tubulin-4A (TUBA4A), which we previously identified as a marker of both migratory periocular NC cells and of CECs (Ramarapu et al., 2026), together with PHH3 and DAPI stain to assess microtubule organization and proliferation. At early stages (day 6), when junctional organization is still immature and nuclei are relatively sparse, TUBA4A displayed a diffuse cytoplasmic distribution (Fig. 7A). This pattern was maintained in PHH3-positive proliferative cells, which were observed at both low and high magnification. Mitotic spindle orientations appeared heterogeneous, with divisions occurring both parallel to the plane of the monolayer and along the apicobasal axis (Fig. 7A, arrows), consistent with the increased variability in nuclear positioning observed at this stage (Fig. 2). As development progressed, TUBA4A expression persisted, and proliferative cells remained detectable, although their frequency decreased by day 10 (Fig. 1D, 7D).

**Figure 7.**
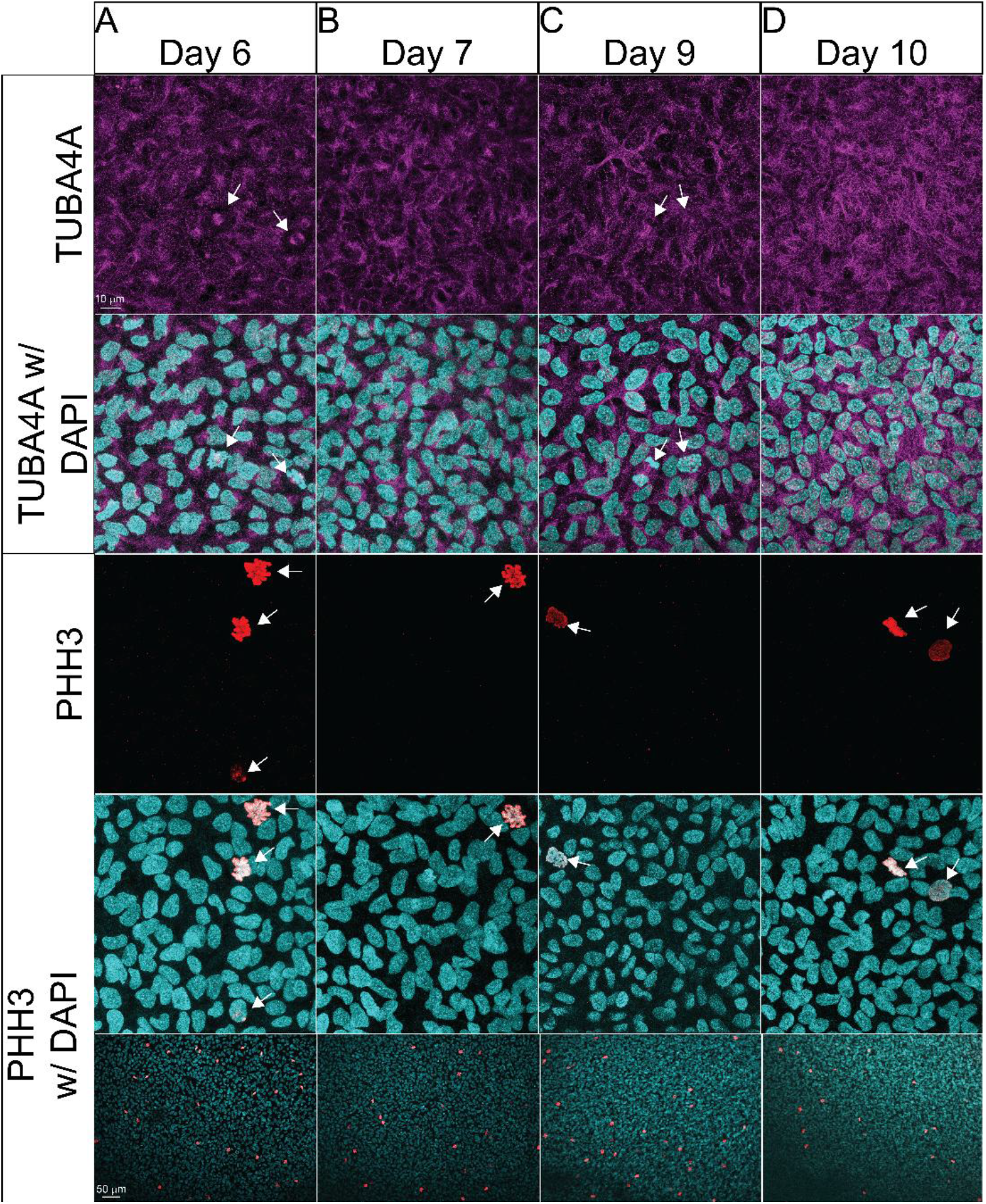
Microtubule organization and proliferative dynamics during early corneal endothelial development. (A-D) Representative confocal images of chick CEC at indicated embryonic stages after IHC for TUBA4A, PHH3 (mitotic cells), and DAPI (nuclei). At day 6 (A), TUBA4A shows diffuse cytoplasmic distribution and is maintained in PHH3-positive cells. Mitotic spindles exhibit variable orientations, including both planar and apicobasal axes (arrows). Proliferative cells are observed at both low and high magnification. As development progresses (B-D), TUBA4A expression persists, while the PHH3-positive cells decreased by day 10, consistent with reduced proliferation. Scale bars: 10 μm in rows 1-4 and 50 μm in bottom row.

To further characterize cytoskeletal organization, we also examined the spatial distribution of TUBA4A across developmental stages. Between days 7 and 8, TUBA4A remained broadly distributed throughout the cytoplasm and near the cell cortex (Fig. 8A,B). By day 11 and later, however, TUBA4A became progressively restricted to the apical region of the cells, forming distinct apical structures and showing reduced signal in basal regions where nuclei are positioned (Fig. 8C-F). This redistribution indicates increasing cytoskeletal polarity during endothelial maturation. High-magnification imaging further revealed the emergence of primary cilia at the apical surface beginning at day 12 (Fig. 8D, white arrows; Fig. S5), consistent with the establishment of apical specialization and functional maturation of the endothelial layer.

**Figure 8.**
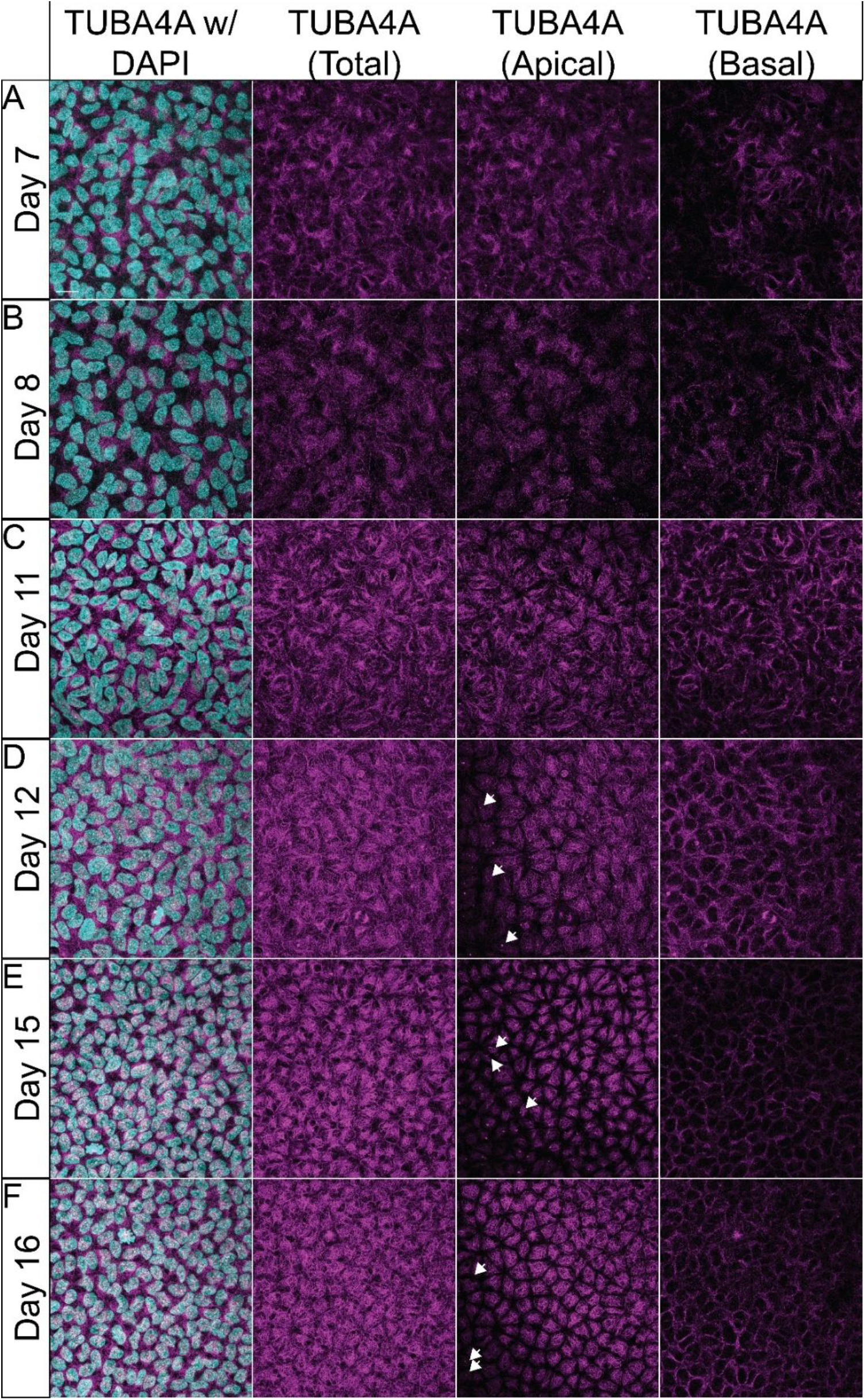
Progressive apical polarization of microtubule organization and emergence of apical specializations. (A-F) Representative confocal images of chick corneal endothelium across developmental stages after IHC for TUBA4A and DAPI (nuclei). At early stages (days 7-8; A, B), TUBA4A is broadly distributed throughout the cytoplasm and near the cell cortex. By day 11 and later (C-F), TUBA4A becomes progressively restricted to the apical region, forming distinct apical structures and showing reduced signal in basal regions where nuclei reside, indicating increasing cytoskeletal polarity. Primary cilia appear to emerge at the apical surface beginning at day 12 (D, white arrows; see also Fig. S5). Scale bars: 10 μm.

## Discussion

The development of a functional corneal endothelium requires coordination between tissue-scale growth and cell-scale organization. By combining quantitative imaging with spatial and topological analysis, our study shows that corneal endothelial maturation is not characterized by large-scale structural transitions or stratification but instead proceeds through a continuous process of geometric equilibration. We propose that temporal decoupling between globe expansion and baseline cell proliferation drives the tissue through distinct mechanical phases, while preserving and progressively refining a low-energy epithelial organization.

A central finding is the persistent dominance of hexagonal topology from the earliest stages examined. In two-dimensional epithelia, hexagonal packing minimizes interfacial free energy and is associated with mechanically stable configurations (Farhadifar et al., 2007; Honda, 1983; Kim et al., 2016; Meineke et al., 2001). In contrast to developmental systems that transition from disorder to order, the chick CEC appears to retain an intrinsic geometric bias from early stages. Maturation therefore reflects progressive reduction of spatial and topological variability rather than *de novo* emergence of order. Consistent with this interpretation, studies in cultured corneal endothelium have shown that mechanical stretch modulates nuclear morphology without disrupting overall geometric constraints (Du et al., 2022).

Our data further suggest that early spatial clustering (Clark-Evans R ≈ 0.78) and elevated z-axis variability represent the final stages of mesenchymal-to-endothelial transition. Migratory cranial NC cells initially exhibit elongated morphologies and overlapping protrusions with limited contact inhibition (Creuzet et al., 2005a; Creuzet et al., 2005b; Ibrahim et al., 2023). The subsequent reduction in Voronoi area variability and rapid smoothing of the z-plane quantitatively captures the establishment of lateral contact inhibition and planar organization. As cells flatten and integrate into the monolayer, inter-nuclear spacing becomes more uniform and the Clark-Evans index increases, reflecting progressive resolution of spatial heterogeneity.

While other cranial NC derivatives also form polarized epithelial-like sheets, such as leptomeninges and odontoblast layers (Arana-Chavez and Massa, 2004; Decimo et al., 2012; Etchevers et al., 2001; Kearns et al., 2023; Santagati and Rijli, 2003; Zhang et al., 2020), the corneal endothelium is uniquely constrained by continuous, multidirectional mechanical strain associated with globe expansion. By contrast, stromal keratocytes derived from later NC migration retain mesenchymal characteristics and are less constrained by strict geometric requirements (Lwigale, 2015). The corneal endothelium must therefore establish epithelial organization while simultaneously accommodating mechanical deformation.

Our cytoskeletal and proliferation analyses suggest mechanisms by which this is achieved. Early in development, proliferative cells exhibit heterogeneous spindle orientations, including divisions both parallel and perpendicular to the plane of the monolayer. Such out-of-plane divisions have been described in other epithelia as a means to accommodate local crowding and integrate new cells into dense tissues (Baena-Lopez et al., 2005; Fouchard et al., 2020; Ragkousi and Gibson, 2014). In this context, apicobasal divisions may transiently displace daughter cells out of the plane, contributing to the localized z-axis variability and spatial clustering observed at early stages. As cells flatten and reintegrate, these perturbations resolve, allowing the tissue to recover a planar configuration.

A second transition occurs around embryonic day 12, marked by a modest increase in z-axis variability, stabilization of nuclear density, and transient deformation of nuclear shape. This stage coincides with a shift from tissue stretching to lateral compaction, suggesting a change in the dominant mechanical regime. Concurrently, we observe progressive maturation of junctional complexes, with NCAD maintained at cell-cell interfaces and ZO-1 transitioning from diffuse cytoplasmic localization to apically enriched tight junctions that align with cortical F-actin. These changes are consistent with the establishment of barrier function and increased mechanical coupling between cells. Notably, corneal thickness and opacity peak at approximately this stage, followed by thinning associated with endothelial pump activation (Conrad et al., 2006).

Cytoskeletal organization appears to play a central role in this transition. Microtubules, initially distributed throughout the cytoplasm, become progressively restricted to the apical domain, forming polarized structures and coinciding with the emergence of primary cilia. This apical polarization likely contributes to junctional maturation and epithelial stability. Previous work has shown that disruption of microtubules can destabilize junctional complexes, leading to redistribution of ZO-1 and breakdown of barrier integrity through RhoA-mediated remodeling of the cortical actin network (Jalimarada et al., 2009). Conversely, stabilization of microtubules can preserve barrier function under stress (Shivanna and Srinivas, 2009). Together with studies of corneal endothelial culture systems (Frausto et al., 2020), these findings support a model in which coordinated microtubule and actin organization underpins junctional integrity and mechanical resilience.

Despite increasing cell density, the corneal endothelium does not exhibit the rise in spatial heterogeneity typically associated with epithelial crowding or “jamming” (Ghosh et al., 2022). Instead, spatial variance continues to decrease as the tissue matures. One possible explanation is that persistent, low-level proliferation acts as a mechanism for stress relaxation. Although the overall proliferation rate declines, variance in mitotic activity remains elevated during intermediate stages, suggesting that cell divisions occur in localized, unsynchronized bursts. This may allow targeted insertion of new cells into high-tension regions, facilitating incremental rearrangement of cell packing without global disruption (Ragkousi and Gibson, 2014).

The increase in the Clark-Evans index from ~0.78 to ~0.91 provides a quantitative signature of this maturation process, reflecting the transition toward a more spatially uniform distribution. By late development, the endothelial lattice closely resembles the organization observed in mature human corneal endothelium, where hexagonality typically ranges from 50-60% (Bourne, 2003; Carlson et al., 1988; Yee et al., 1985).

Taken together, our findings support a model in which corneal endothelial development is driven by continuous geometric refinement under mechanical constraint. By integrating imaging with quantitative spatial analysis, this study provides a framework for linking tissue-scale growth, cytoskeletal organization, and epithelial topology. More broadly, these principles are likely to apply to other systems in which epithelial tissues must maintain structural order while adapting to dynamic mechanical environments.

## Materials and Methods

### Chicken eggs, incubation, and dissection

Fertilized chicken eggs were obtained from the University of California Davis Hopkins Avian Facility and incubated at 37°C for developmental stages encompassing embryonic days 6 and 16. Following incubation, tissues were dissected, cleared of yolk and blood using Ringer’s solution and fixed based on developmental stage. For early stages (day 6-10), the anterior section of both globes in each embryo were dissected along the corneal/limbal margin with the remainder of the globe attached to the whole embryo. For later stages, the globes were dissected from the embryos, the eyelids and lens were removed, following which the cornea was trimmed along the limbal margin. Surrounding tissue such as the iris was removed if damaged to minimize blood cell autofluorescence and pigment staining on the corneal endothelium. 3-4 radial incisions were made, if necessary, along the corneal margin to minimize curvature. Tissues were fixed at 4% paraformaldehyde (PFA) for 20 minutes at room temperature.

### Macroscopic globe metrics

To determine globe metrics, images of globes were acquired immediately post dissection (Table 1). Due to fragility of the early embryonic eye, day 6-10 globes were imaged while still attached to the embryo, angled perpendicularly to the camera. For later stages, globes were imaged post-dissection. All macroscopic images were acquired with a calibration scale bar placed at the exact same focal plane as the globe to ensure accurate spatial scaling. Images were processed using ImageJ (FIJI) (Schneider et al., 2012). The perimeter of each globe was manually traced using the polygonal outliner tool, and the “Measure” function was utilized to extract absolute globe surface area (mm^2^) and Feret’s diameter. These macroscopic metrics were subsequently exported to R for integration with the cellular-level morphometric datasets.

### Fluorescence Imaging and Image Processing - 2D Imaging pipeline

For 2D spatial and geometric analysis, images were acquired using a Zeiss Axio Imager.M2 with Apotome capability at 10X, 20X, or 63X magnification. For high-resolution 3D topological mapping, flatmounted images were acquired on a Leica SP8 STED 3X confocal microscope controlled by Leica LAS X software. Images were captured using a high numerical aperture 100x/1.4 oil immersion objective (HC PL APO CS2). Z-stacks were acquired with a precise 0.3 μm step size. To maximize signal-to-noise ratio and optical resolution, all confocal images were deconvolved using Huygens Batch Feeder version 25.10 (Scientific Volume Imaging, The Netherlands) utilizing a sample template deconvolution strategy.

### 2D and 3D Nuclear Segmentation

Automated 2D nuclear instance segmentation was performed on the Zeiss-acquired images using a generalist Cellpose-SAM model (Pachitariu et al., 2025). A 63X magnification image was cropped to a 100 µm by 100 µm section and segmentation was performed using a flow threshold of 0.2 and a cell probability threshold of 1.0. Nuclear count was measured as the number of objects in the mask and the nuclear mask was exported to CellProfiler (Stirling et al., 2021) for downstream analysis. Nuclei at the image border were excluded using the FilterObjects module, and the MeasureObjectSizeShape module was used to characterize all morphological measurements. Number of embryos and images used for analysis in Table 2.

For 3D nuclear data, the deconvolved Leica Z-stacks were imported into Imaris (version 10.2). The Imaris “Surfaces” function was utilized to automatically generate high-fidelity 3D reconstructions of the DAPI-stained nuclei. From these surfaces, 3D centroid coordinates (X, Y, Z) were extracted and exported for Z-MAD mathematical modeling. Imaris was also utilized to generate 3D surface rendering images isolating the nuclear geometries, which were exported as high-resolution TIFFs for figure assembly.

### Immunohistochemistry and flat mounting

Following fixation, corneal samples were washed with 1X TBS containing 0.5% Triton X-100 (TBST) with calcium. To minimize non-specific binding, samples were incubated in a blocking buffer consisting of PBST with 10% donkey serum for 1 to 7 days at 4 °C. Primary antibodies (including markers for mitotic spindles, TUBA4A, and active proliferation, PHH3) as detailed in Table 3 were diluted in blocking buffer and incubated with the tissue for approximately 72 hours at 4 °C. After primary incubation, samples were thoroughly washed with TBST+Ca^2+^ and subsequently incubated with AlexaFluor secondary antibodies (1:500) and a DAPI nuclear counterstain (1:500) in blocking buffer for up to 24 hours at 4 °C. AlexaFluor 568 Phalloidin (1:100) was added for corresponding samples with secondary antibodies. Following secondary incubation, tissues were washed in TBST and post-fixed in 4% PFA for 1 hour at room temperature to stabilize the fluorescent complexes. For imaging, the corneas were carefully flat mounted radially on glass slides using Fluoromount-G and cover slipped.

**Table 3.** Antibodies utilized in study.

| Antibody Target | Antigen | Dilution | Species Reactivity | Host/Isotype | Source |
| --- | --- | --- | --- | --- | --- |
| Tubulin alpha-4A (TUBA4A) | Full-length protein | 1:250 | Human | Mouse/IgG2b | ThermoFisher |
| Phosphorylated Histone H3 (PHH3) | KLH-conjugated linear peptide corresponding to human Histone H3 (Ser10) | 1:500 | Human | Rat/IgG2a | Sigma-Aldrich |
| N-Cadherin (NCAD) | Amino acids 308-597 | 1:5 | Chick, Human, Mouse, *Xenopus* | Rat/IgG2a | DSHB |
| ZO-1 | Amino acids 1446-1748 from recombinant human ZO1 protein | 1:250 | Human, Mouse, Rat, Pig, Rabbit, Canine, Chick, Bovine, Goat | Rabbit/IgG | Proteintech |

### Proliferation Quantification

Automated quantification of proliferation was performed on the Zeiss-acquired images at 63X magnification taken at random locations within the central cornea in a PHH3-blind manner. Images were cropped to remove the scale bar, and nuclear count was determined as described above. The PHH3 count for each image was then determined manually, and the proliferation rate was calculated as the ration of PHH3 positive cells to overall nuclear count for each image.

### Voronoi Tessellation and Geometric Analysis

To mathematically model the topological architecture of the endothelial monolayer, Voronoi tessellations were generated from the 2D nuclear centroid coordinates using the *deldir* package in R. To eliminate geometric distortion caused by the boundary limits of the field of view, strict edge-correction was applied. For the remaining true-interior cells, the number of polygon sides (including hexagonality) and the intercellular area variance were calculated, with the latter quantified as the Coefficient of Variation (CV) of the interior Voronoi areas.

### 2D Spatial Statistics and point pattern analysis

Macroscopic spacing and organization of the monolayer were evaluated using the spatstat package in R (Baddeley A, 2015). Inter-nuclear spacing was evaluated by calculating the absolute Nearest Neighbor Distance (NND) for every nucleus, aggregating the mean and standard deviation (SD) of the NND per image. To determine the degree of spatial regularity independent of global tissue density, the Clark-Evans Aggregation Index (R) was calculated as the ratio of the observed mean NND to the expected mean NND of a random Poisson distribution of identical density. R = 1 indicates complete spatial randomness, R < 1 indicates clustering, and R > 1 indicates regularized/equidistant spacing.

### 3D Spatial Regression and Monolayer Topology

To accurately quantify the apical-basal flatness of the monolayer and eliminate artifacts from tightly apposed stromal keratocytes, a robust morpho-spatial filter was utilized. For each 3D image, a 2D spatial Loess regression (Z ~ X * Y, span = 0.4) was fitted to the nuclear centroids to approximate the curved centerline of the endothelial sheet. The absolute residual distance of each nucleus from this theoretical sheet (ΔZ) was calculated. The structural “flatness” and topology of the endothelial monolayer were then quantified for each image as the Median Absolute Deviation (MAD) of the ΔZ values.

### Statistical Analysis

All statistical testing was performed in R. To ensure measurement precision, macroscopic globe area was independently quantified by two investigators; inter-rater reliability was confirmed via Pearson correlation, and averaged values were used for subsequent analysis (Fig. S1, Table S1). For 2D image-level aggregate metrics (e.g., nuclear Density, Clark-Evans R, Area CV, NND, and Mean PHH3+ Index), developmental transitions were evaluated using non-parametric Wilcoxon rank-sum tests, treating independent fields-of-view as replicates but averaging image-level data to the embryo level to conservatively avoid pseudoreplication. Developmental shifts in the synchronization of proliferation were evaluated by testing the variance of the PHH3+ index using the Fligner-Killeen test.

To rigorously evaluate global 3D topological flatness (Z-MAD) across developmental stages, differences were analyzed using Linear Mixed-Effects Models (LMM) via the *lme4* package over N=3 biological replicates per time point (Bates et al., 2015). Developmental Age was treated as a fixed effect, while the specific embryo of origin was modeled as a random intercept to account for intra-sample correlation (Formula ~ Age + (1|Embryo_ID)). Significance thresholds were set at α = 0.05. Statistical analyses Rmarkdown code, input data and output of all comparisons performed are available on github: https://github.com/luckycharms10/CEC-Architecture-Study.

## Acknowledgements

The authors acknowledge the numerous members of the Comparative Ophthalmology and Vision Sciences Laboratory and to the members of the Rogers Laboratory at UC Davis, UC Davis School of Veterinary Medicine for their contributions throughout the study period. Special appreciation to the UC Davis Hopkins Avian Facility and the UC Davis Advanced Imaging Facility for their contiued support.

## Competing interests

The authors declare that they have no competing interests.

## Funding

Financial support for this work was provided by grants from: the National Institutes of Health including the National Eye Institute (R01 EY016134 and R01 EY036440) to SMT; National Institute of Dental and Craniofacial Research (NIDCR) R03 DE032047-01, National Science Foundation (NSF) 2143217, and Hypothesis Fund Catalyst Grant to CDR; and the UC Davis Vision Science T32 (NEI-EY015387) to support RR.

## Data and resource availability

Rmarkdown file of statistical analysis, input data is available on GitHub (https://github.com/luckycharms10/CEC-Architecture-Study) and raw imaging data is available upon request.

## Notes

https://github.com/luckycharms10/CEC-Architecture-Study

### Competing Interest Statement

The authors have declared no competing interest.

## References

Arana-Chavez, V. E. and Massa, L. F. (2004). Odontoblasts: the cells forming and maintaining dentine. Int J Biochem Cell Biol 36, 1367–73.

Babushkina, A. and Lwigale, P. (2020). Periocular neural crest cell differentiation into corneal endothelium is influenced by signals in the nascent corneal environment. Dev Biol 465, 119–129.

Baddeley A R. E.,, Turner R. (2015). Spatial Point Patterns: Methodology and Applications with R. Chapman and Hall/CRC Press, London.

Baena-Lopez, L. A., Baonza, A. and Garcia-Bellido, A. (2005). The orientation of cell divisions determines the shape of Drosophila organs. Curr Biol 15, 1640–4.

Bates, D., Mächler, M., Bolker, B. and Walker, S. (2015). Fitting Linear Mixed-Effects Models Using lme4. Journal of Statistical Software 67, 1 – 48.

Bourne, W. M. (2003). Biology of the corneal endothelium in health and disease. Eye (Lond) 17, 912–8.

Campbell, H. K., Maiers, J. L. and DeMali, K. A. (2017). Interplay between tight junctions & adherens junctions. Exp Cell Res 358, 39–44.

Carlson, K. H., Bourne, W. M., McLaren, J. W. and Brubaker, R. F. (1988). Variations in human corneal endothelial cell morphology and permeability to fluorescein with age. Exp Eye Res 47, 27–41.

Conrad, A. H., Zhang, Y., Walker, A. R., Olberding, L. A., Hanzlick, A., Zimmer, A. J., Morffi, R. and Conrad, G. W. (2006). Thyroxine affects expression of KSPG-related genes, the carbonic anhydrase II gene, and KS sulfation in the embryonic chicken cornea. Invest Ophthalmol Vis Sci 47, 120–32.

Coulombre, A. J. (1957). The role of intraocular pressure in the development of the chick eye. II. Control of corneal size. AMA Arch Ophthalmol 57, 250–3.

Creuzet, S., Couly, G. and Le Douarin, N. M. (2005a). Patterning the neural crest derivatives during development of the vertebrate head: insights from avian studies. J Anat 207, 447–59.

Creuzet, S., Vincent, C. and Couly, G. (2005b). Neural crest derivatives in ocular and periocular structures. Int J Dev Biol 49, 161–71.

Cvekl, A. and Tamm, E. R. (2004). Anterior eye development and ocular mesenchyme: new insights from mouse models and human diseases. Bioessays 26, 374–86.

Decimo, I., Fumagalli, G., Berton, V., Krampera, M. and Bifari, F. (2012). Meninges: from protective membrane to stem cell niche. Am J Stem Cells 1, 92–105.

Du, R., Li, D., Huang, Y., Xiao, H., Xue, J., Ji, J., Feng, Y. and Fan, Y. (2022). Effect of mechanical stretching and substrate stiffness on the morphology, cytoskeleton and nuclear shape of corneal endothelial cells. Medicine in Novel Technology and Devices 16, 100180.

Etchevers, H. C., Vincent, C., Le Douarin, N. M. and Couly, G. F. (2001). The cephalic neural crest provides pericytes and smooth muscle cells to all blood vessels of the face and forebrain. Development 128, 1059–68.

Farhadifar, R., Roper, J. C., Aigouy, B., Eaton, S. and Julicher, F. (2007). The influence of cell mechanics, cell-cell interactions, and proliferation on epithelial packing. Curr Biol 17, 2095–104.

Fouchard, J., Wyatt, T. P. J., Proag, A., Lisica, A., Khalilgharibi, N., Recho, P., Suzanne, M., Kabla, A. and Charras, G. (2020). Curling of epithelial monolayers reveals coupling between active bending and tissue tension. Proc Natl Acad Sci U S A 117, 9377–9383.

Frausto, R. F., Swamy, V. S., Peh, G. S. L., Boere, P. M., Hanser, E. M., Chung, D. D., George, B. L., Morselli, M., Kao, L., Azimov, R. et al. (2020). Phenotypic and functional characterization of corneal endothelial cells during in vitro expansion. Sci Rep 10, 7402.

Gage, P. J., Rhoades, W., Prucka, S. K. and Hjalt, T. (2005). Fate maps of neural crest and mesoderm in the mammalian eye. Invest Ophthalmol Vis Sci 46, 4200–8.

Ghosh, B., Nishida, K., Chandrala, L., Mahmud, S., Thapa, S., Swaby, C., Chen, S., Khosla, A. A., Katz, J. and Sidhaye, V. K. (2022). Epithelial plasticity in COPD results in cellular unjamming due to an increase in polymerized actin. J Cell Sci 135.

Goldminz, D., Vlodavsky, I., Johnson, L. K. and Gospodarowicz, D. (1979). Contact inhibition and the regulation of endocytosis in the corneal endothelium: correlation with a restricted surface receptor lateral mobility and the appearance of a fibronectin meshwork. Exp Eye Res 29, 331–51.

Harrison, T. A., He, Z., Boggs, K., Thuret, G., Liu, H. X. and Defoe, D. M. (2016). Corneal endothelial cells possess an elaborate multipolar shape to maximize the basolateral to apical membrane area. Mol Vis 22, 31–9.

Hay, E. D. and Revel, J. P. (1969). Fine structure of the developing avian cornea. Monogr Dev Biol 1, 1–144.

Honda, H. (1983). Geometrical models for cells in tissues. Int Rev Cytol 81, 191–248.

Ibrahim, N., Hifny, A., Elhanbaly, R., El-Desoky, S. M. M. and Gaber, W. (2023). Morphogenetic events influencing corneal maturation, development, and transparency: Light and electron microscopic study. Microsc Res Tech 86, 539–555.

Jalimarada, S. S., Shivanna, M., Kini, V., Mehta, D. and Srinivas, S. P. (2009). Microtubule disassembly breaks down the barrier integrity of corneal endothelium. Exp Eye Res 89, 333–43.

Kearns, N. A., Iatrou, A., Flood, D. J., De Tissera, S., Mullaney, Z. M., Xu, J., Gaiteri, C., Bennett, D. A. and Wang, Y. (2023). Dissecting the human leptomeninges at single-cell resolution. Nat Commun 14, 7036.

Kim, S., Cassidy, J. J., Yang, B., Carthew, R. W. and Hilgenfeldt, S. (2016). Hexagonal Patterning of the Insect Compound Eye: Facet Area Variation, Defects, and Disorder. Biophys J 111, 2735–2746.

Lindner, T., Klose, R., Streckenbach, F., Stahnke, T., Hadlich, S., Kuhn, J. P., Guthoff, R. F., Wree, A., Neumann, A. M., Frank, M. et al. (2017). Morphologic and biometric evaluation of chick embryo eyes in ovo using 7 Tesla MRI. Sci Rep 7, 2647.

Lwigale, P. Y. (2015). Corneal Development: Different Cells from a Common Progenitor. Prog Mol Biol Transl Sci 134, 43–59.

Maurice, D. M. (1957). The structure and transparency of the cornea. J Physiol 136, 263–86.

Meineke, F. A., Potten, C. S. and Loeffler, M. (2001). Cell migration and organization in the intestinal crypt using a lattice-free model. Cell Prolif 34, 253–66.

Pachitariu, M., Rariden, M. and Stringer, C. (2025). Cellpose-SAM: superhuman generalization for cellular segmentation. bioRxiv, 2025.04.28.651001.

Petroll, W. M., Hsu, J. K., Bean, J., Cavanagh, H. D. and Jester, J. V. (1999). The spatial organization of apical junctional complex-associated proteins in feline and human corneal endothelium. Curr Eye Res 18, 10–9.

Ragkousi, K. and Gibson, M. C. (2014). Cell division and the maintenance of epithelial order. J Cell Biol 207, 181–8.

Ramarapu, R., Stoehr, W. R., Miesen, M., Border, S., Thomasy, S. and Rogers, C. D. (2026). A molecular and spatial resource defining tubulin isotype organization during corneal development. bioRxiv, 2026.02.19.706651.

Roy, O., Leclerc, V. B., Bourget, J. M., Theriault, M. and Proulx, S. (2015). Understanding the process of corneal endothelial morphological change in vitro. Invest Ophthalmol Vis Sci 56, 1228–37.

Santagati, F. and Rijli, F. M. (2003). Cranial neural crest and the building of the vertebrate head. Nat Rev Neurosci 4, 806–18.

Schmid, G. F. (2003). Axial and peripheral eye length measured with optical low coherence reflectometry. J Biomed Opt 8, 655–62.

Schneider, C. A., Rasband, W. S. and Eliceiri, K. W. (2012). NIH Image to ImageJ: 25 years of image analysis. Nat Methods 9, 671–5.

Shivanna, M. and Srinivas, S. P. (2009). Microtubule stabilization opposes the (TNF-alpha)-induced loss in the barrier integrity of corneal endothelium. Exp Eye Res 89, 950–9.

Steffek, A. J., Mujwid, D. K. and Johnston, M. C. (1979). Scanning electron microscopy (SEM) of cranial neural crest migration in chick embryos. Birth Defects Orig Artic Ser 15, 11–21.

Stirling, D. R., Swain-Bowden, M. J., Lucas, A. M., Carpenter, A. E., Cimini, B. A. and Goodman, A. (2021). CellProfiler 4: improvements in speed, utility and usability. BMC Bioinformatics 22, 433.

Yee, R. W., Matsuda, M., Schultz, R. O. and Edelhauser, H. F. (1985). Changes in the normal corneal endothelial cellular pattern as a function of age. Curr Eye Res 4, 671–8.

Zhang, M., Zhang, X., Luo, J., Yan, R., Niibe, K., Egusa, H., Zhang, Z., Xie, M. and Jiang, X. (2020). Investigate the Odontogenic Differentiation and Dentin-Pulp Tissue Regeneration Potential of Neural Crest Cells. Front Bioeng Biotechnol 8, 475.

