## Supplementary Table and Figures for "Quantitative imaging of corneal endothelial development reveals dynamic but resilient monolayer"

**Table S1. Daily changes in globe size paired with nuclei density**.

**Table of Daily Morphometric Rate of Change**

Allometric scaling of Globe Area, Endothelial Nuclei Density

| | Globe Area Expansion | | CEC Nuclei | |
|---|---|---|---|---|
| **Age** | **Globe Growth %** | **Globe Sig** | **Density Growth %** | **Density Sig** |
| Day 6 | - | - | - | - |
| Day 7 | 135.89 | **** | 10.19 | ns |
| Day 8 | 70.43 | **** | 0.71 | ns |
| Day 9 | 20.35 | * | 4.9 | ns |
| Day 10 | 25.47 | *** | 22.67 | **** |
| Day 11 | 8.06 | ns | 0.98 | ns |
| Day 12 | 7.88 | ns | 20.16 | *** |
| Day 13 | 5.18 | ns | 7.66 | ns |
| Day 14 | 10.35 | ** | 12.88 | * |
| Day 15 | 5.36 | ns | -1.13 | ns |
| Day 16 | 11.99 | *** | -4.52 | ns |

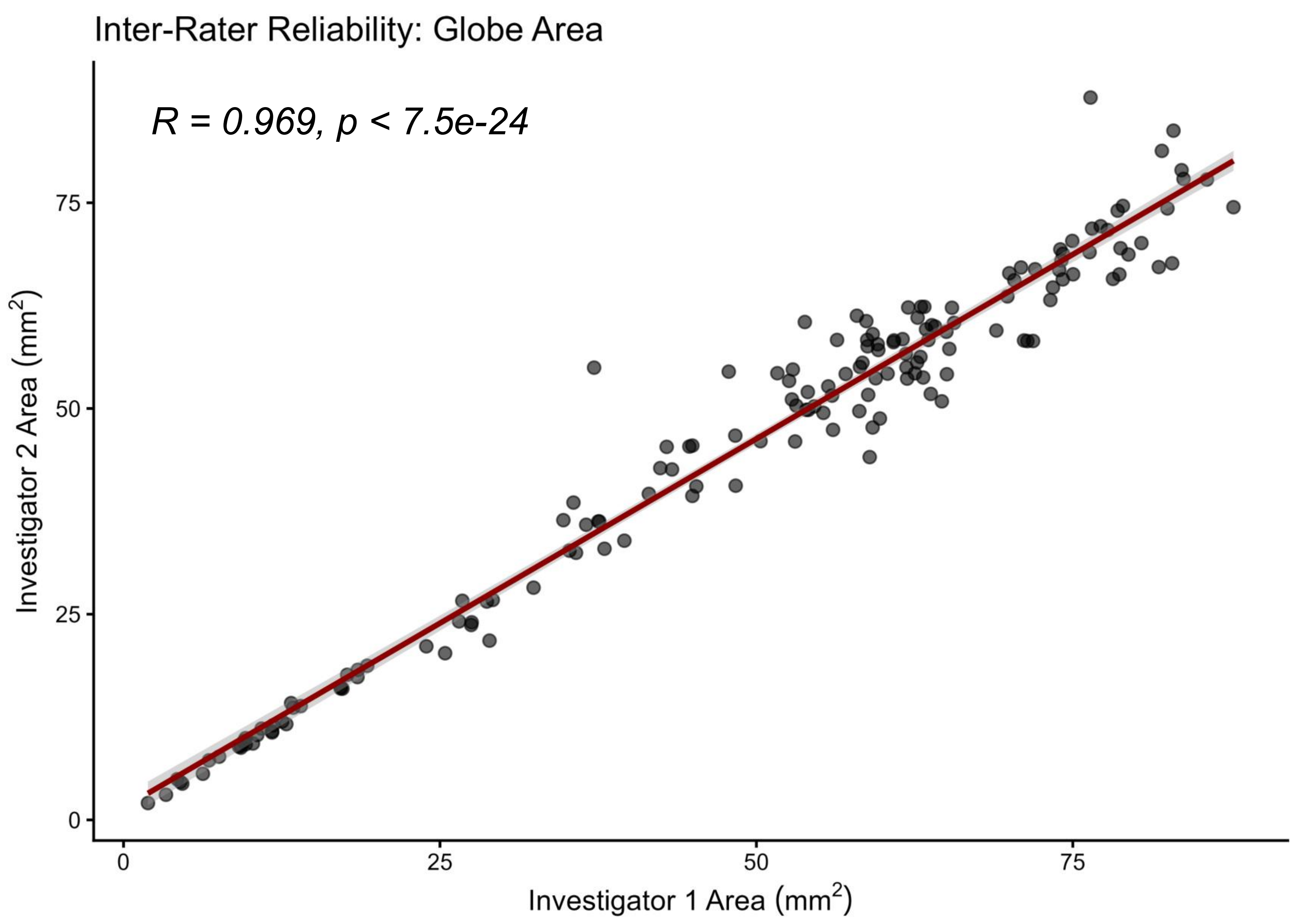


**Fig. S1. Scatter plot of inter-rater reliability for imageJ measures of globe size**. Globe area (selected measure) for matched samples plotted for each investigator. Pearson correlation statistical output provides high confidence for measurement precision.

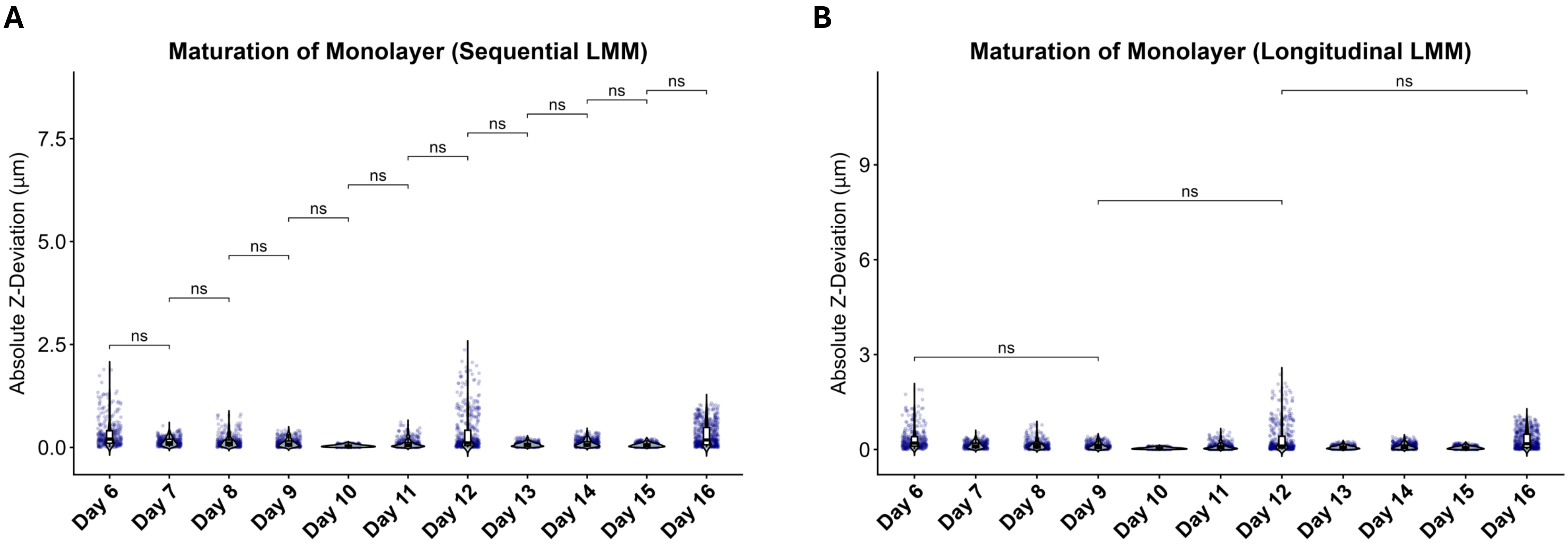


**Fig. S2. Violin plots showing distribution of absolute Z-deviation of 3D nuclear seed positions across the developmental timeline.** (A) Plot demonstrating sequential linear mixed model statistics. (B) Plot showing longitudinal linear mixed model statistics. Ns, non-significant.

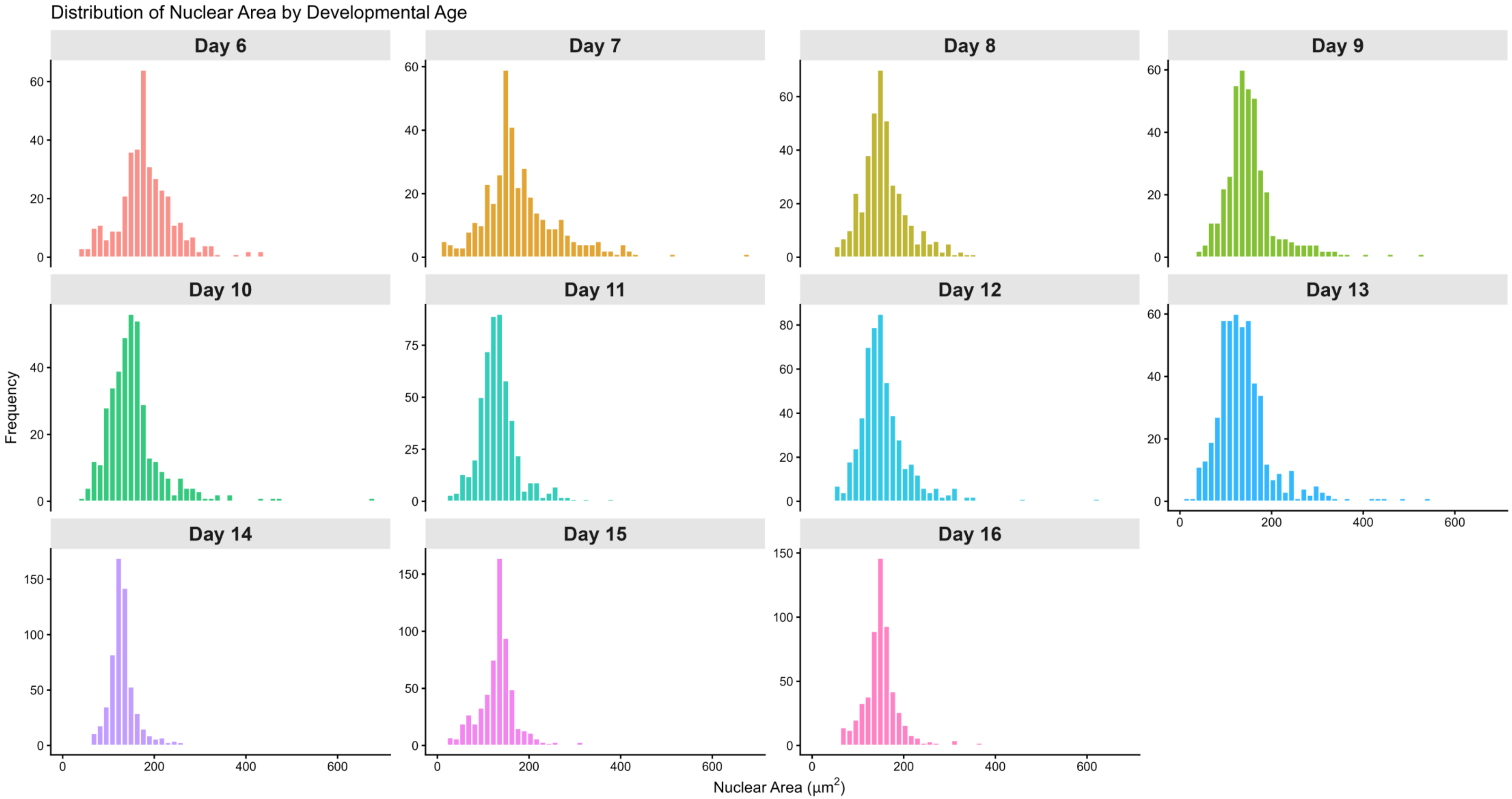


**Fig. S3. Histograms demonstrating distribution of 3d nuclei area by stage.** Bins utilized are size 40.

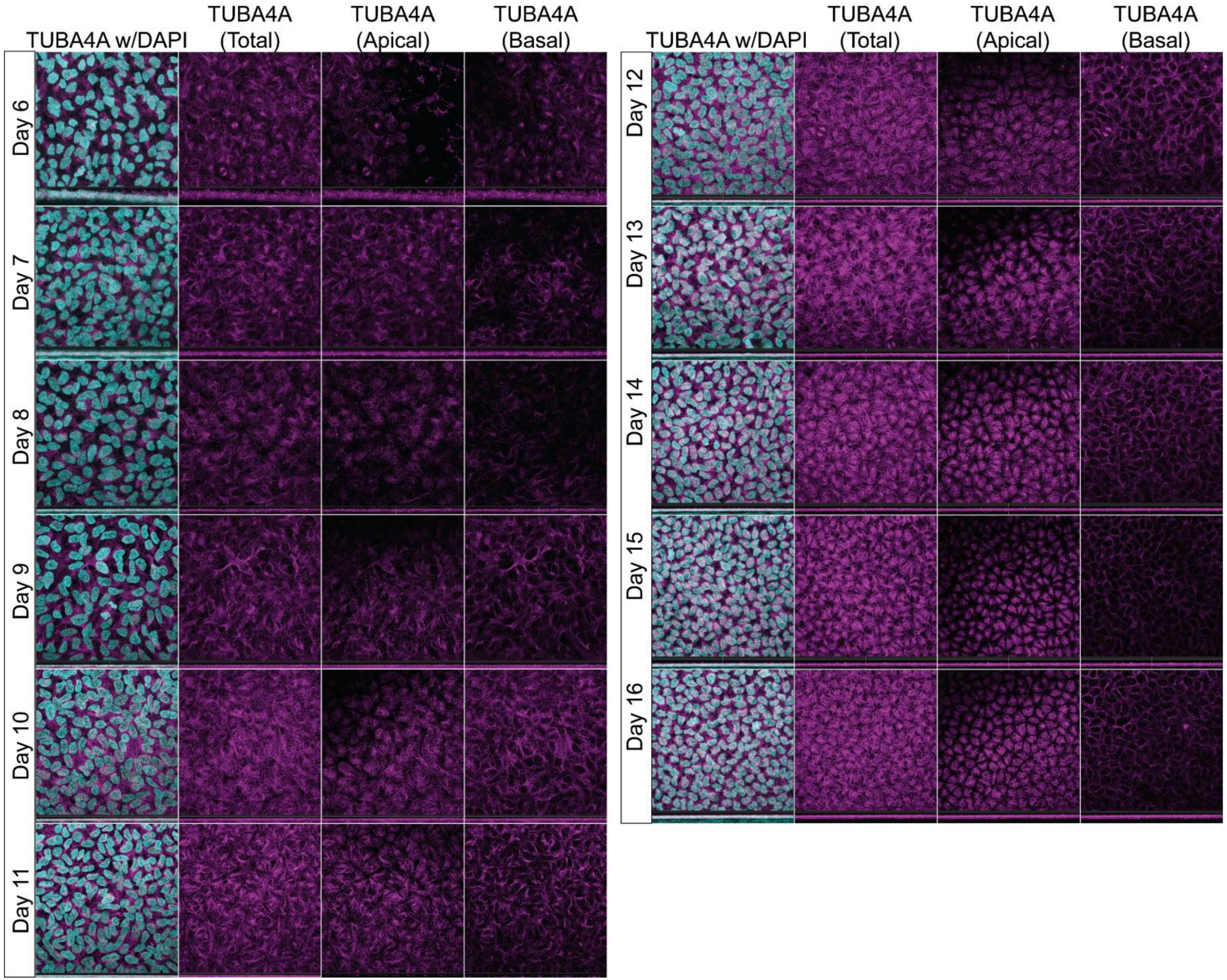


**Fig. S4. TUBA4A apicobasal localization from day 6-16.** IHC for TUBA4A with DAPI stain on corneal flatmounts from day 6-16. Column 1: Maximum intensity projection (MIP) image showing both DAPI (cyan) and TUBA4A (magenta). Column 2: MIP of total CEC TUBA4A signal. Column 3: Apical MIP showing changing TUBA4A patterning in apical CEC. Column 4: MIP of basal TUBA4A signal showing perimembranous distribution.

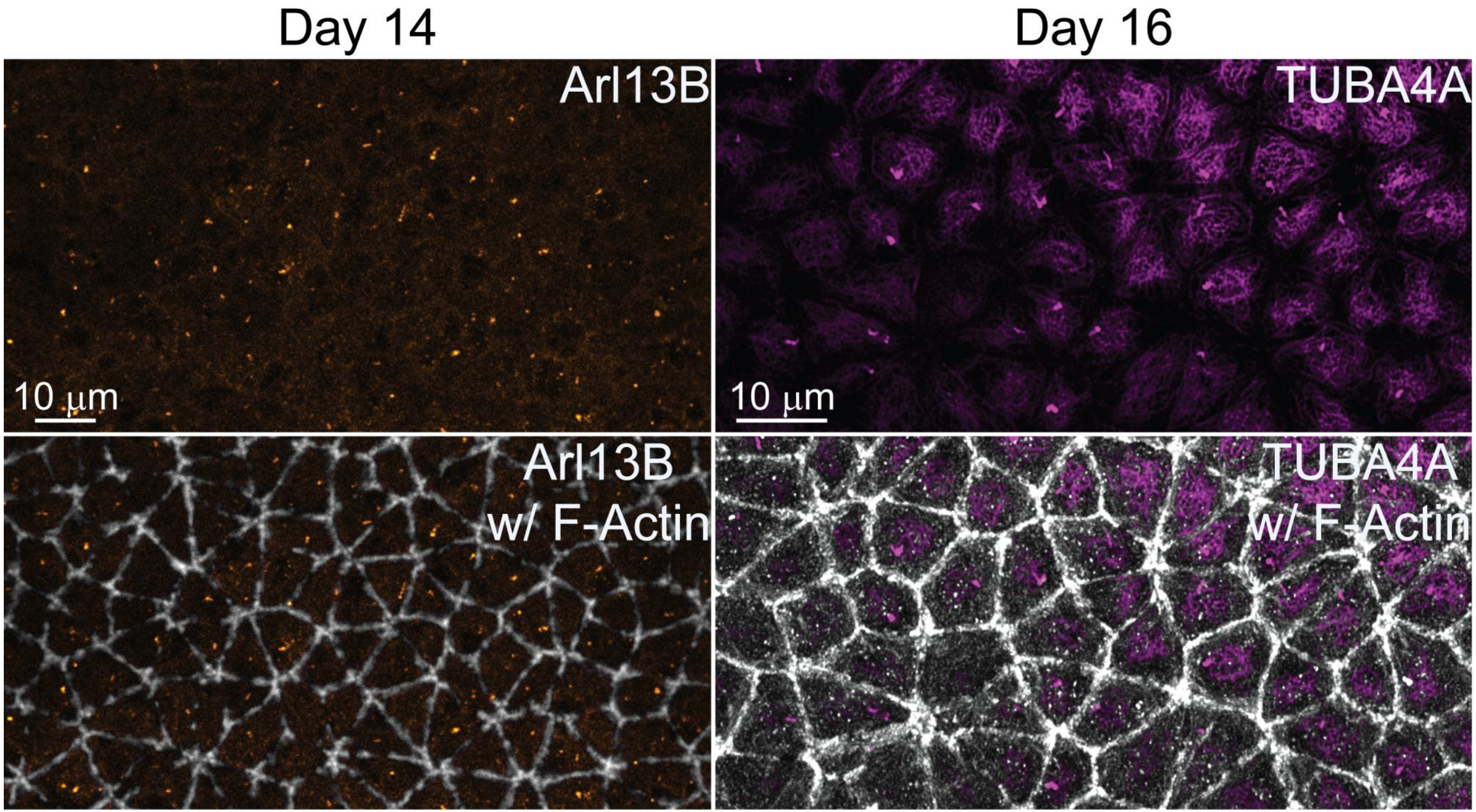


**Fig. S5. Arl13B and TUBA4A apical localization at days 14 and 16.** IHC for Arl13B (orange) and TUBA4A (magenta) with F-actin stain on corneal flatmounts at days 14 and 16.
